## Supplementary File for "Choosing the Best Route: Comparative Optimization of Wheat Transformation Methods for Improving Yield by Targeting *TaARE1-D* with CRISPR/Cas9"

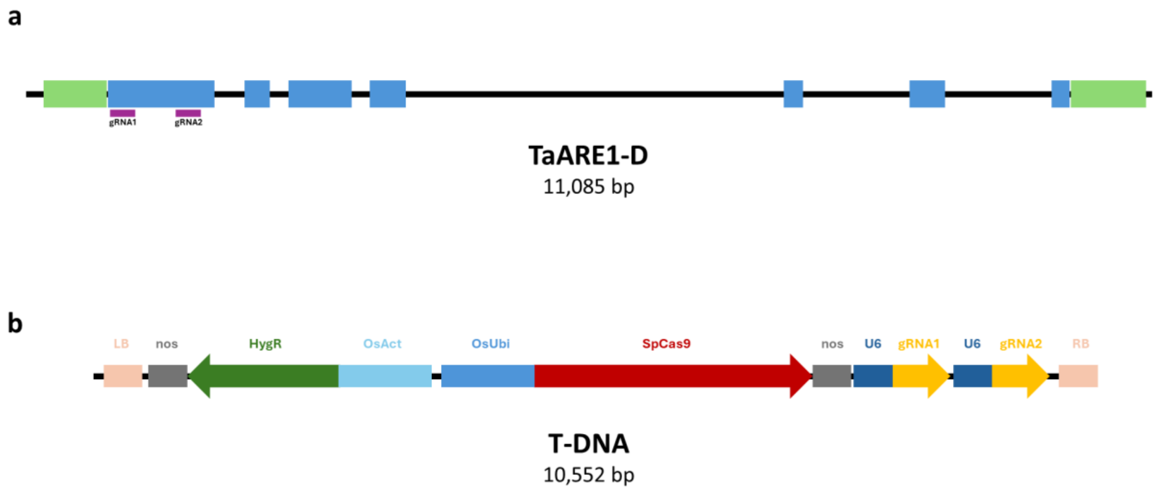


**Supplementary Figure 1.** Schematic representation of the TaARE1-D gene structure in *T. aestivum* and the CRISPR/Cas9 constructs used for editing. Untranslated regions (UTRs, green) and exons (blue) are shown as rectangles, with gRNA target sites indicated (a). The CRISPR/Cas9 plasmid contained *OsAct:HygR:nos* for selection, SpCas9 driven by the *OsUbi* promoter, and *TaU6* as the pol III promoter for expression of both gRNAs.

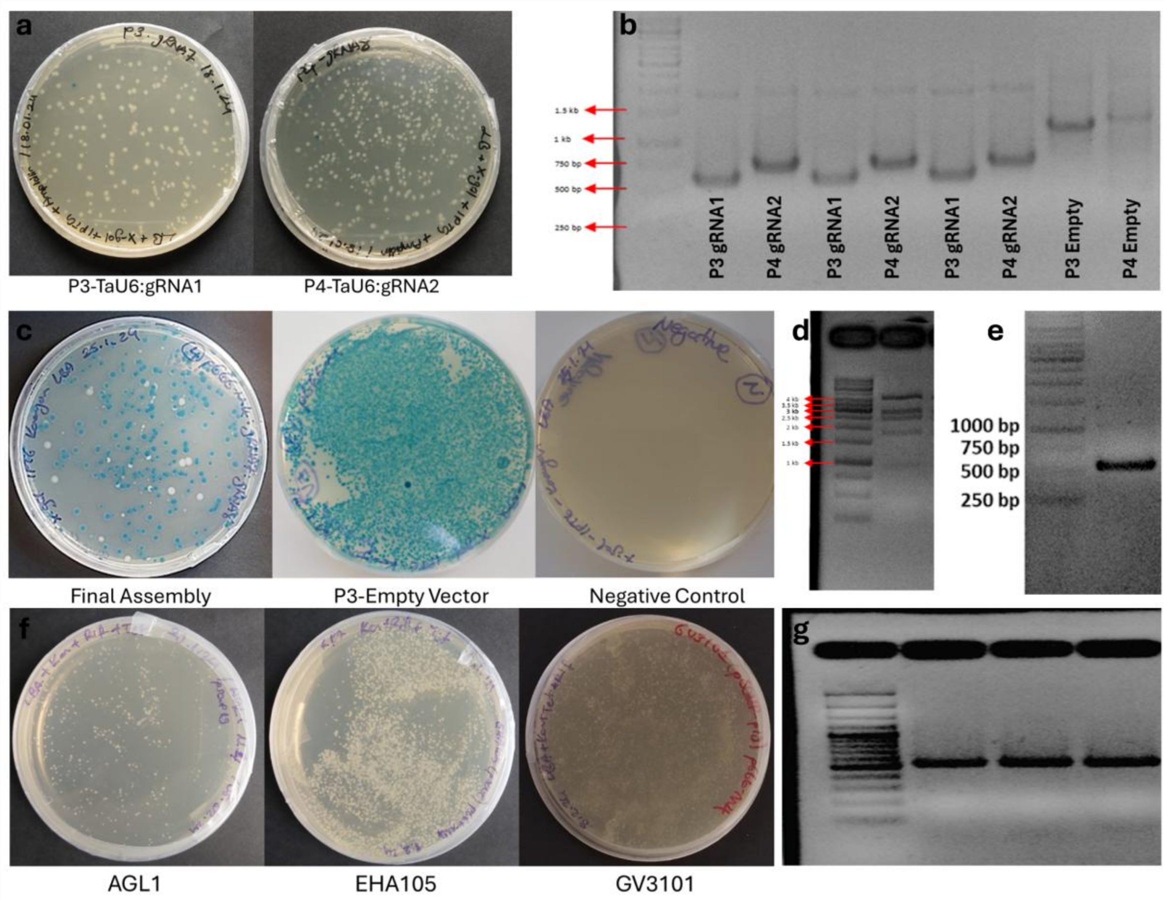


**Supplementary Figure 2.** Cloning and verification of CRISPR/Cas9 plasmids targeting *TaARE1-D*. The P3:TaU6:gRNA1 and P4:TaU6:gRNA2 entry plasmids were transformed into *E. coli* DH5α and subjected to blue/white screening (a). Plasmids were isolated from selected white colonies and verified by PCR using TaU6 Sequencing-F and L440 primers, producing shorter amplicons in P3 and P4 compared to empty vectors due to the removal of *lacZ* (b). The final Golden Gate reaction was performed for construct assembly and confirmed by blue/white screening (c), restriction digestion (d), and PCR with gRNA1 sense and gRNA2 antisense primers (e). Verified plasmids were subsequently transferred into *Agrobacterium tumefaciens* strains AGL1, EHA105, and GV3101 by electroporation. Colonies were selected on rifampicin, tetracycline, and kanamycin (f) and confirmed by colony PCR using gRNA1 sense and gRNA2 antisense primers (g)

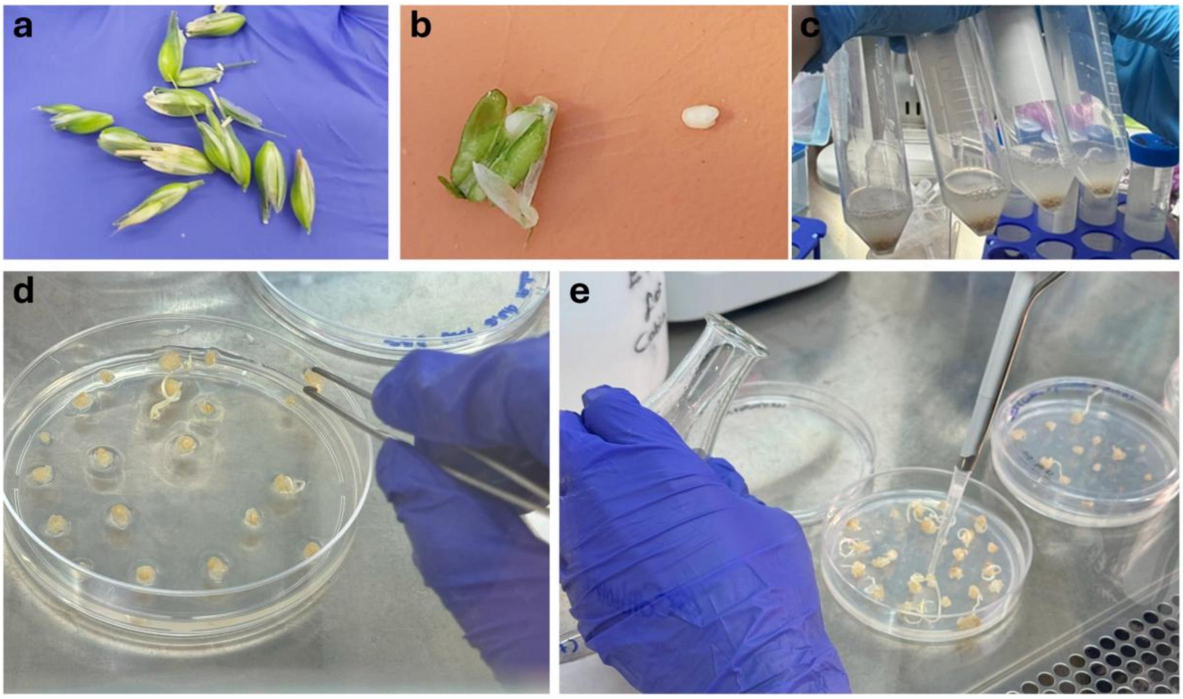


**Supplementary Figure 3.** Transformation of immature embryos and callus derived from mature embryos. Immature embryos were excised from spikelets 2–3 weeks after anthesis (a, b) and subjected to transformation with *Agrobacterium tumefaciens* strains AGL1, EHA105, and GV3101 (c). Callus was induced from mature embryos placed on culture medium (d) and subsequently used for transformation, during which the callus surface was covered with *Agrobacterium* suspension (e).

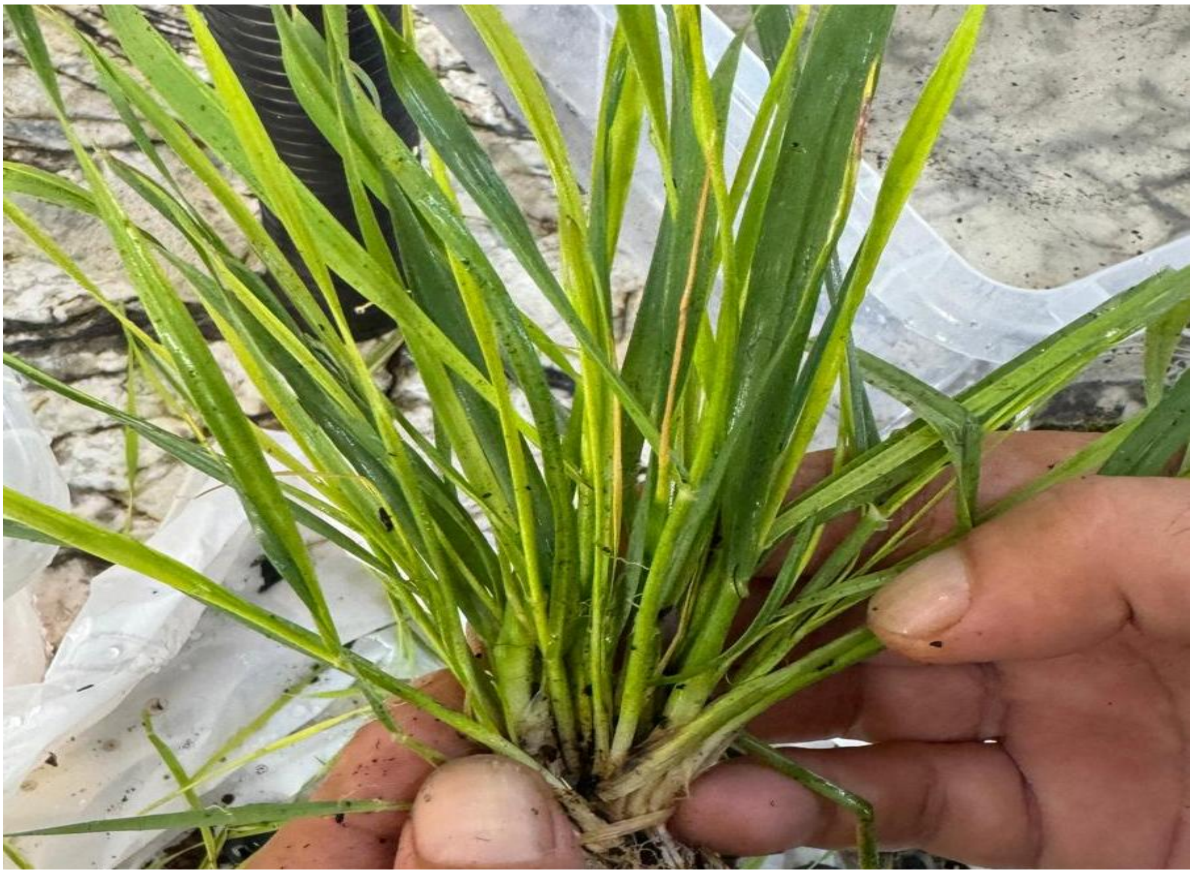

**Supplementary Figure 4.** Over-tillering observed in wheat plants transformed by the *in planta* method three weeks after Agrobacterium injection.

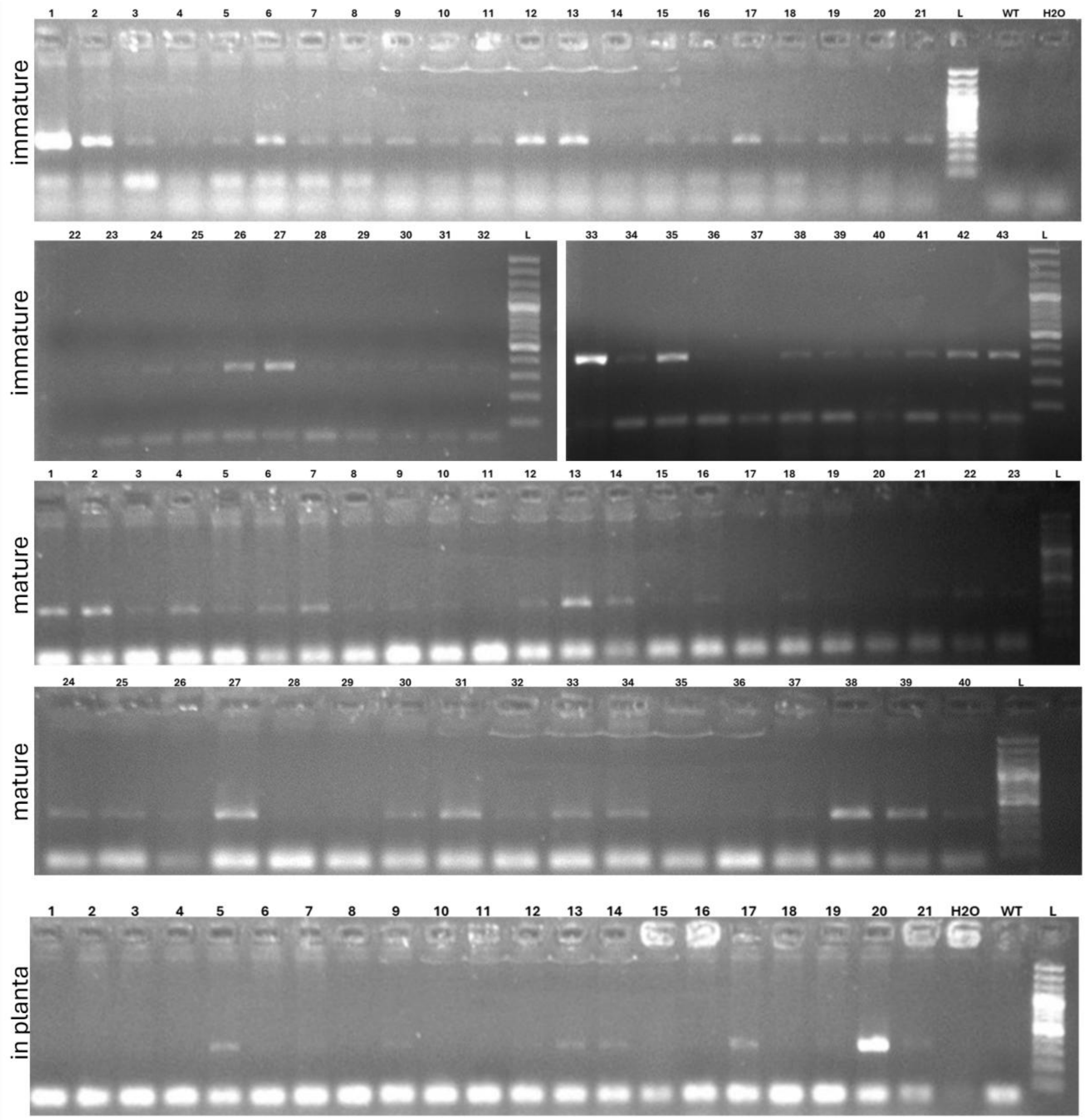


**Supplementary Figure 5.** Transgene screening of regenerated wheat plants derived from immature embryo, mature embryo callus, and *in planta* transformations. Plants obtained from immature and mature embryos were screened after hygromycin selection, whereas *in planta*-transformed plants were screened 4–5 weeks after Agrobacterium AGL1 injection.

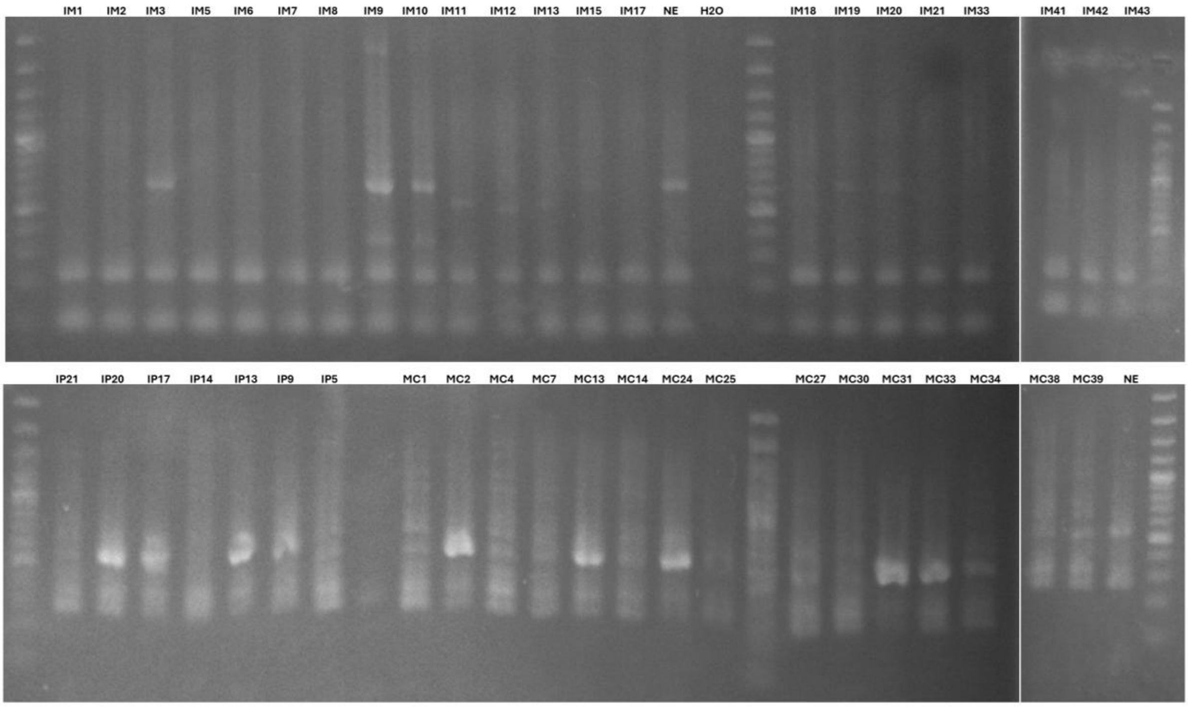


**Supplementary Figure 6.** Screening of regenerated transgenic wheat plants using ACT-PCR with ACT-F/R primers. Amplification products on the gel indicate the presence of the non-edited *TaARE1-D* allele, whereas the absence of a band indicates at least one mutation induced by CRISPR/Cas9 due to primer mismatch at the gRNA1 target site.

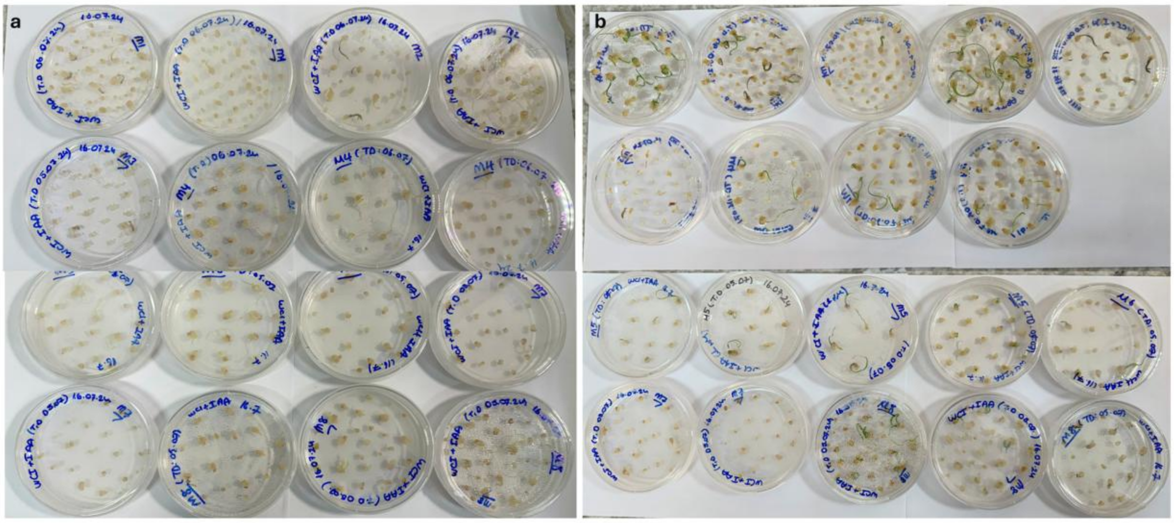


**Supplementary Figure 7.** Callus after Agrobacterium-mediated transformation (a) and regenerated calli one week after transfer to selection medium supplemented with 0.2 mg/l IAA (b).

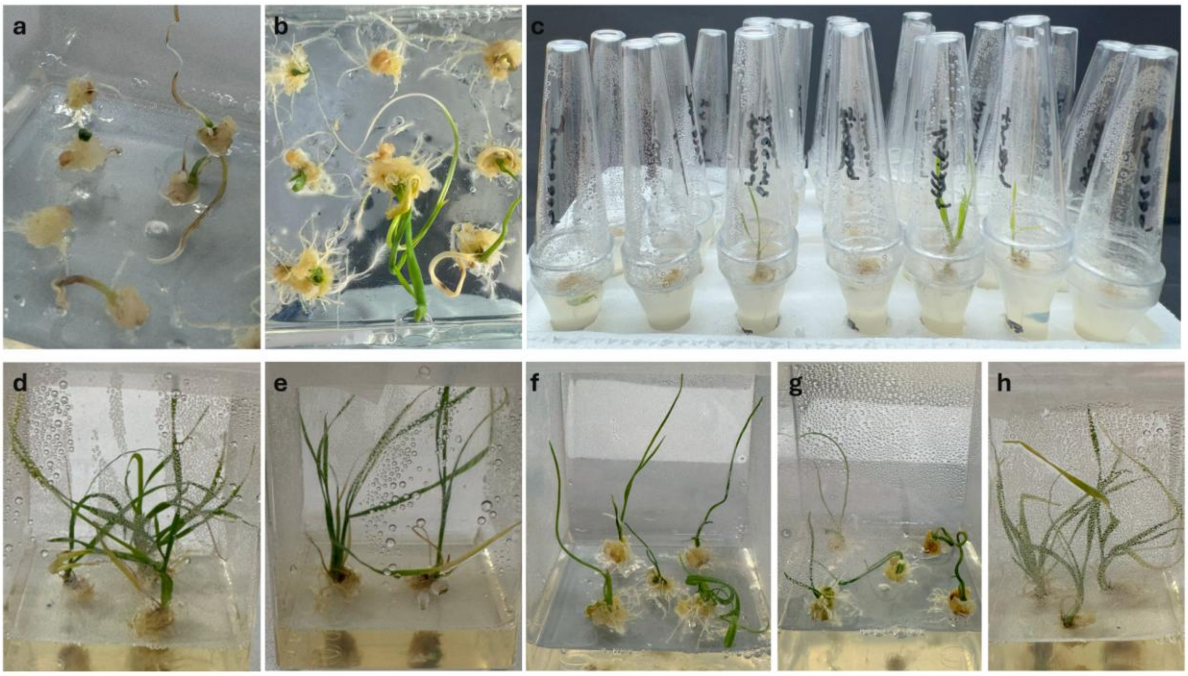


**Supplementary Figure 8.** Regenerated calli derived from transformed immature embryos (a–c), and regenerated plants after two months on selection medium, ready for acclimatization (d–h).

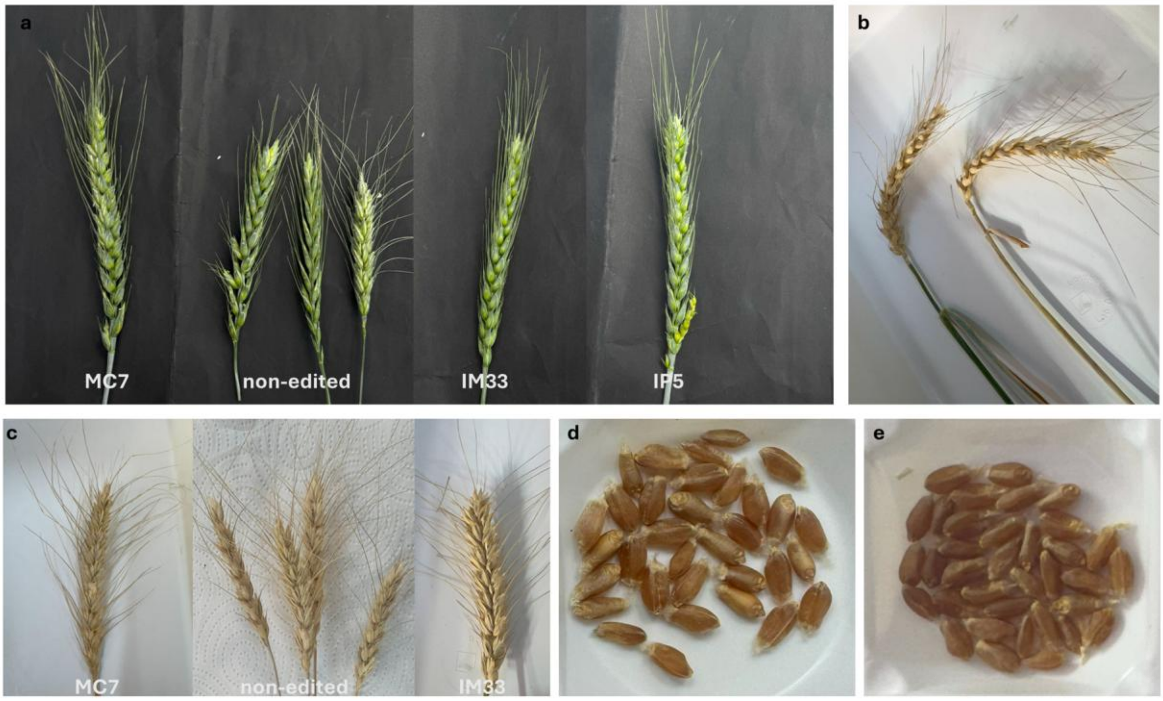


**Supplementary Figure 9.** Spike morphology of NE, IM33, MC7, and IP5 (a); stay-green phenotype of IM33 (left) showing green sap despite dried spikes compared to NE (right) (b); spikes of NE, MC7, and IM33 ready for harvesting (c); and collected seeds from edited lines (d–e).

**Supplementary Table 1.** Sequences of gRNAs and primers that are used in the study

| Oligonucleotide Name | Sequences | Purpose |
|---|---|---|
| gRNA1-sense | CTTGTGCGGTCTGGTTCGCCGTAG | gRNA1 cloning |
| gRNA1-antisense | AAACCTACGGCGAACCAGACCGCA | gRNA1 cloning |
| gRNA2-sense | CTTGCCAGATAACGGATGATCCGA | gRNA2 cloning |
| gRNA2-antisense | AAACTCGGATCATCCGTTATCTGG | gRNA2 cloning |
| TaU6 Sequencing-F | TAGGAGGGAATCGAACTAGGAATATTG | Plasmid Verification |
| L440 | AGCGAGTCAGTGAGCGAG | Plasmid Verification |
| ACT-F | TGCGGTCTGGTTCGCCGTAG | Screening |
| ACT-R | AGCAATGTTGCACAGTAATAAGG | Screening |
| TaARE1D-F | CCCGGAAGATGTGGTCTTCG | Screening |
| TaARE1D-R | TTAGTGCTAGAAGTACCGATTTCG | Screening |

**Supplementary Table 2.** Immature embryo transformation and average survival rates under hygromycin selection, evaluated across different *Agrobacterium* strains, optical densities (OD), acetosyringone concentrations, and incubation times ($p = 0.01$).

| Strain | OD | Acetosyringone (μM) | Time of Duration (mins) | Average (%) | SD | Group |
|---|---|---|---|---|---|---|
| AGL1 | 0.8 | 150 | 15 | 61.67 | 0.0523 | a |
| EHA105 | 0.8 | 150 | 15 | 37.30 | 0.0404 | b |
| GV3101 | 0.8 | 150 | 15 | 10.42 | 0.0779 | c |
| AGL1 | 0.6 | 150 | 15 | 48.90 | 0.0455 | b |
| AGL1 | 0.8 | 150 | 15 | 75.64 | 0.0090 | a |
| AGL1 | 1.0 | 150 | 15 | 24.52 | 0.0351 | c |
| AGL1 | 0.8 | 100 | 15 | 41.51 | 0.0117 | b |
| AGL1 | 0.8 | 150 | 15 | 68.38 | 0.0120 | a |
| AGL1 | 0.8 | 200 | 15 | 20.20 | 0.1449 | b |
| AGL1 | 0.8 | 150 | 10 | 27.47 | 0.0323 | c |
| AGL1 | 0.8 | 150 | 15 | 63.95 | 0.0210 | a |
| AGL1 | 0.8 | 150 | 20 | 51.10 | 0.0455 | b |

**Supplementary Table 3.** Callus transformation derived from mature embryos and average survival rates under hygromycin selection, assessed according to *Agrobacterium* strain, optical density (OD), acetosyringone concentration, and incubation time ($p = 0.01$).

| Strain | OD | Acetosyringone (μM) | Time of Duration (mins) | Average (%) | SD | Group |
|---|---|---|---|---|---|---|
| AGL1 | 0.8 | 150 | 15 | 51.68 | 0.0650 | a |
| EHA105 | 0.8 | 150 | 15 | 26.59 | 0.0498 | b |
| GV3101 | 0.8 | 150 | 15 | 12.27 | 0.0902 | b |
| AGL1 | 0.6 | 150 | 15 | 49.11 | 0.0665 | a |
| AGL1 | 0.8 | 150 | 15 | 57.51 | 0.0315 | a |
| AGL1 | 1.0 | 150 | 15 | 25.71 | 0.0404 | b |
| AGL1 | 0.8 | 100 | 15 | 48.93 | 0.0814 | a |
| AGL1 | 0.8 | 150 | 15 | 57.91 | 0.0315 | a |
| AGL1 | 0.8 | 200 | 15 | 12.64 | 0.0388 | b |
| AGL1 | 0.8 | 150 | 10 | 73.93 | 0.0664 | a |
| AGL1 | 0.8 | 150 | 15 | 63.50 | 0.0696 | a |
| AGL1 | 0.8 | 150 | 20 | 27.68 | 0.0237 | b |

**Supplementary Table 4.** Comparison of regeneration efficiency between 0.2 mg/l IAA and 1 mg/l zeatin in callus derived from immature and mature embryos ($p = 0.01$).

| Explant | Hormone | Average Regeneration Rate (%) | SD | Group |
|---|---|---|---|---|
| Immature | Zeatin | 22.77 | 0.0522 | b |
| Mature | Zeatin | 24.55 | 0.0420 | b |
| Immature | IAA | 50.70 | 0.0732 | a |
| Mature | IAA | 47.94 | 0.0490 | a |

**Supplementary Table 5.** *In planta* transformation and average survival rate under hygromycin selection according to Agrobacterium strain, optical density (OD), acetosyringone concentration, and co-cultivation time ($p = 0.01$).

| Strain | OD | Acetosyringone (μM) | Time of Duration (days) | Average (%) | SD | Group |
|---|---|---|---|---|---|---|
| AGL1 | 0.8 | 150 | 3 | 30.42 | 0.0278 | a |
| EHA105 | 0.8 | 150 | 3 | 10.69 | 0.0316 | b |
| GV3101 | 0.8 | 150 | 3 | 2.22 | 0.0314 | b |
| AGL1 | 0.6 | 150 | 3 | 7.01 | 0.0546 | c |
| AGL1 | 0.8 | 150 | 3 | 23.17 | 0.246 | b |
| AGL1 | 1.0 | 150 | 3 | 39.19 | 0.0492 | a |
| AGL1 | 0.8 | 100 | 3 | 32.54 | 0.0876 | ab |
| AGL1 | 0.8 | 150 | 3 | 36.51 | 0.0449 | a |
| AGL1 | 0.8 | 200 | 3 | 15.28 | 0.0335 | b |
| AGL1 | 0.8 | 150 | 2 | 38.73 | 0.0339 | a |
| AGL1 | 0.8 | 150 | 3 | 24.52 | 0.0351 | b |
| AGL1 | 0.8 | 150 | 4 | 27.14 | 0.0487 | ab |

**Supplementary Table 6.** Phenotypic observation of edited and non-edited lines and their statistical groups (*p=0.01*)

| line | Number of Grain Per Spike | Spike Length (cm) | Grain Length (cm) | Thousand Grain Weight (g) |
|---|---|---|---|---|
| NE | 52.00 | 8.96 | 0.49 | 39.11 |
| NE | 54.00 | 8.98 | 0.54 | 39.56 |
| NE | 56.00 | 9.22 | 0.62 | 40.98 |
| NE | 56.00 | 9.36 | 0.64 | 41.12 |
| NE | 57.00 | 9.48 | 0.68 | 40.98 |
| NE | 58.00 | 9.66 | 0.69 | 41.14 |
| NE | 59.00 | 9.68 | 0.71 | 40.07 |
| NE | 59.00 | 9.72 | 0.74 | 39.99 |
| NE | 62.00 | 9.77 | 0.78 | 40.11 |
| NE | 62.00 | 9.88 | 0.79 | 42.11 |
| **NE Average** | 57.5 | 9.471 | 0.669 | 40.517 |
| **NE SD** | 2.899843256 | 0.297823926 | 0.088534534 | 0.812225338 |
| **group** | b | b | b | b |
| IM33 | 64.00 | 9.87 | 0.64 | 41.02 |
| IM33 | 59.00 | 9.93 | 0.76 | 41.89 |
| IM33 | 66.00 | 10.11 | 0.77 | 42.12 |
| IM33 | 67.00 | 10.22 | 0.78 | 42.44 |
| IM33 | 68.00 | 10.28 | 0.78 | 42.66 |
| IM33 | 61.00 | 10.36 | 0.82 | 44.43 |
| IM33 | 69.00 | 10.55 | 0.84 | 44.98 |
| IM33 | 65.00 | 10.69 | 0.85 | 45.55 |
| IM33 | 66.00 | 10.96 | 0.89 | 45.97 |
| IM33 | 64.00 | 11.02 | 0.96 | 46.66 |
| **IM33 Average** | 65 | 10.33 | 0.791888889 | 43.772 |
| **IM33 SD** | 3.055050463 | 0.335062183 | 0.068010529 | 1.693559332 |
| **group** | a | a | a | a |
| MC7 | 61.00 | 9.89 | 0.61 | 39.98 |
| MC7 | 61.00 | 9.98 | 0.64 | 42.96 |
| MC7 | 61.00 | 10.07 | 0.70 | 42.98 |
| MC7 | 63.00 | 10.09 | 0.72 | 43.56 |
| MC7 | 65.00 | 10.09 | 0.74 | 43.98 |
| MC7 | 66.00 | 10.09 | 0.77 | 44.09 |
| MC7 | 66.00 | 10.13 | 0.78 | 44.12 |
| MC7 | 66.00 | 10.24 | 0.78 | 44.42 |
| MC7 | 67.00 | 10.59 | 0.80 | 44.56 |
| MC7 | 69.00 | 10.62 | 0.93 | 45.05 |
| **MC7 Average** | 64.50 | 10.18 | 0.746 | 43.57 |
| **MC7 SD** | 2.692582404 | 0.2298456 | 0.085807284 | 1.350644291 |
| **group** | a | a | ab | a |
| IP5 | 61 | 10.00 | 0.65 | 38.99 |
| IP5 | 61 | 10.03 | 0.66 | 39.96 |
| IP5 | 62 | 10.05 | 0.70 | 42.16 |
| IP5 | 62 | 10.11 | 0.75 | 42.56 |
| IP5 | 62 | 10.13 | 0.75 | 44.12 |
| IP5 | 64 | 10.14 | 0.76 | 44.42 |
| IP5 | 64 | 10.16 | 0.77 | 44.52 |
| IP5 | 65 | 10.19 | 0.79 | 44.56 |
| IP5 | 65 | 10.56 | 0.84 | 44.68 |
| IP5 | 66 | 10.72 | 0.88 | 44.96 |
| **IP5 Average** | 63.2 | 10.209 | 0.7544 | 43.093 |
| **IP5 SD** | 1.720465053 | 0.225585904 | 0.069560334 | 2.020227957 |
| **group** | a | a | ab | a |
